# Distinguishing the activity of adjacent somatosensory nuclei within the brainstem using 3T fMRI

**DOI:** 10.1101/2024.11.27.625699

**Authors:** Paige Howell, Ingrid Odermatt, Olivia Harrison, Finn Rabe, Sarah Meissner, Patrick Freund, Nicole Wenderoth, Sanne Kikkert

## Abstract

Experimental evidence in animal models indicates that the brainstem plays a major role in sensory modulation. However, mapping functional activity within the human brainstem presents many methodological challenges. These constraints have deterred essential research into human sensory brainstem processing. Here, using a 3T fMRI sequence optimised for the brainstem, combined with uni- and multivariate analysis approaches, we investigated the extent to which functional activity of neighbouring somatosensory nuclei can be delineated in the brainstem, thalamus and primary somatosensory cortex (S1). Whilst traditional univariate approaches offered limited differentiation between adjacent hand and face activation in the brainstem, multivariate classification enabled above-chance decoding of these activity patterns across S1, the thalamus, and the brainstem. Our findings establish a robust methodological approach to explore signal processing within the brainstem and across the entire somatosensory stream. This is a fundamental step towards broadening our understanding of somatosensory processing within humans and determining what changes in sensory integration may occur in clinical populations following sensory deprivation.

## 1. Introduction

Accurately localising touch and proprioception is vital for daily functioning. If our brain were to mislocate a hand touch as being on the face, it would cause disoriented movements, making simple tasks like eating or dressing nearly impossible. The processing of afferent inputs is organised in a somatotopic fashion along the somatosensory processing stream, which facilitates the accurate interpretation of afferent signals and coordination of movements. This map-like organisation showcases a point-to-point correspondence between spatial activity patterns in the central nervous system (CNS) and body parts. For example, tactile and proprioceptive signals of the hands enter the CNS via the dorsal horn and then travel through the spinal cord before reaching the first CNS relay station in the ipsilateral cuneate nucleus of the medulla (Figure 1). From the cuneate nucleus, the signals are projected to the contralateral ventroposterior lateral (VPL) nucleus of the thalamus and finally to the hand area of the primary somatosensory cortex and higher-order cortical areas (Purves, 2019). Conversely, afferent signals from the face synapse adjacently in the ipsilateral trigeminal nuclei before reaching the contralateral ventroposterior medial (VPM) nucleus of the thalamus and, finally, the face area of the primary somatosensory cortex and higher-order cortical regions (Craven, 2011; Purves, 2019).

**Figure 1.**
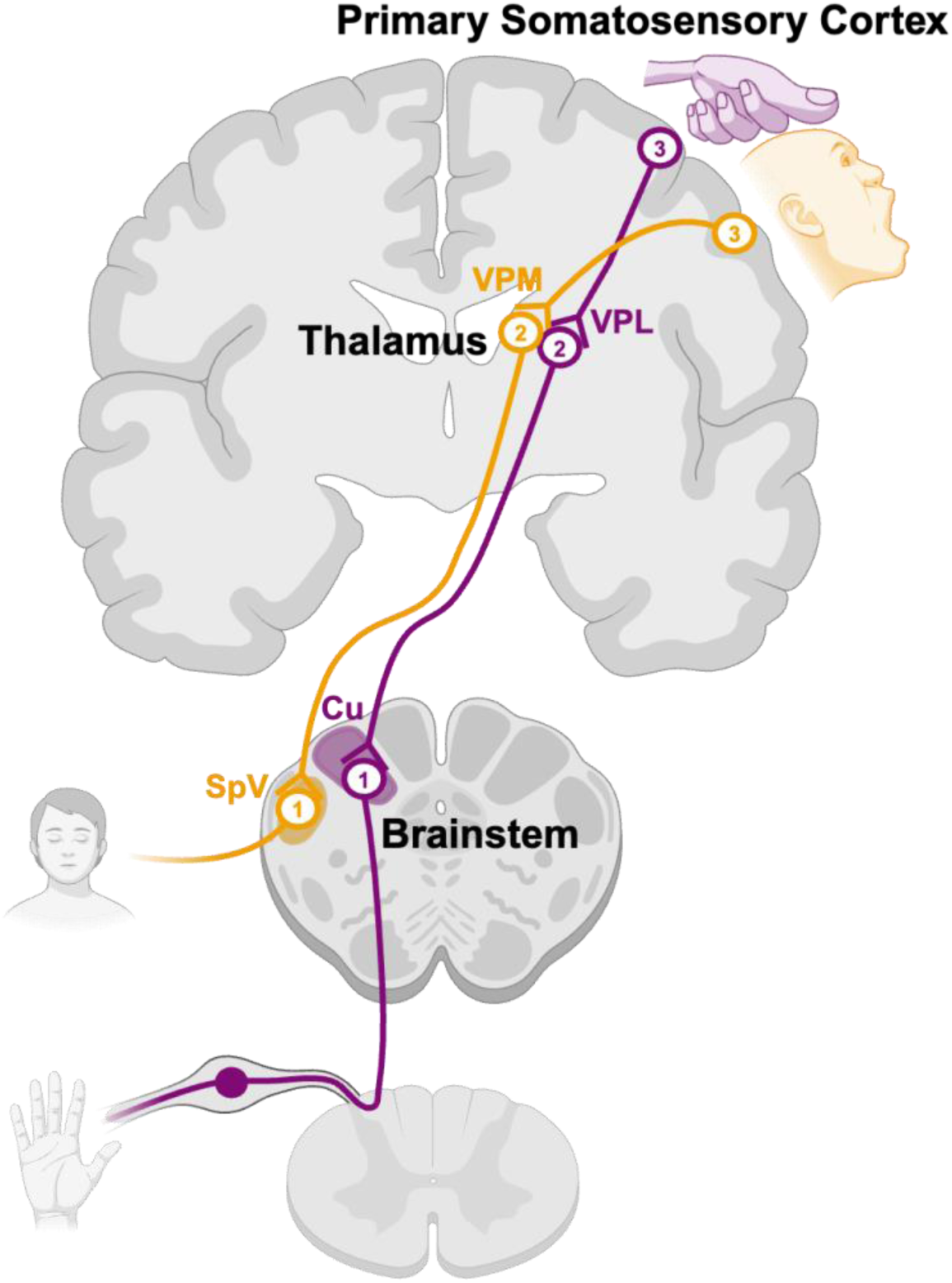
Processing of tactile and proprioceptive inputs across the central nervous system. Tactile and proprioceptive sensations from the hand (purple) ascend via the ipsilateral dorsal column of the spinal cord, reaching the cuneate (Cu) nucleus in the brainstem (1; purple). From there, signals are relayed to the contralateral ventroposterior lateral (VPL) nucleus of the thalamus (2; purple) and then to the primary somatosensory cortex (3; purple). Conversely, inputs from the face (yellow) are relayed to the ipsilateral trigeminal (SpV) nucleus in the brainstem (1; yellow) before projecting to the contralateral ventroposterior medial (VPM) nucleus of the thalamus (2; yellow) and then to the primary somatosensory cortex (3; yellow). Figure created with BioRender.com.

The brainstem somatosensory nuclei have been termed the somatosensory gateway to the brain (Versteeg et al., 2021). Studies in animal models have indicated that they play a critical role in sensory modulation by attenuating, enhancing or rerouting incoming signals (Kubota et al., 2024; Versteeg, Rosenow et al., 2021; Versteeg, Chowdhury et al., 2021; Jain et al., 2000; Keine et al., 2016). For instance, a recent study in non-human primates demonstrated that somatosensory signals are attenuated in the cuneate nucleus during voluntary movement as compared to passive movement (Kubota et al., 2024). Additionally, non-human primate models of spinal cord injury have identified these brainstem nuclei as key sites for driving upstream plasticity, where reorganisation between inputs from the adjacent cuneate and trigeminal nuclei leads to the cortical expansion of the face representations in the primary somatosensory cortex (S1) (Halder et al., 2018; Jain et al., 2000; Kaas et al., 1999; Kambi et al., 2014). This plasticity has been linked to both recovery of function and the development of maladaptive circuitry that relates to neuropathic pain (Jutzeler, Curt, & Kramer, 2015; Zaaimi, Edgley, Soteropoulos & Baker, 2012; Zörner et al., 2014). Therefore, investigating the brainstem is not only crucial for a complete systems-level understanding of somatosensory processing but may also provide insight into the driving forces of adaptive plasticity in clinical cohorts with impaired somatosensation. However, despite its importance, our knowledge of brainstem somatosensory processing in humans remains limited, as it is extremely challenging to explore human brainstem nuclei in vivo. Importantly, the relationship between somatosensory organisation in the brainstem and the plasticity of cortical representations remains unexplored in humans. Here, as a fundamental step towards imaging functionally distinct somatosensory processing pathways in humans, we investigated the extent to which we can delineate the activity of neighbouring somatosensory nuclei across the brainstem, thalamus and S1, using 3T functional Magnetic Resonance Imaging (fMRI).

fMRI offers a non-invasive solution to probe brainstem function in vivo. However, while it is regularly used to assess somatotopic organisation and processing in cortical areas, specifically S1 (Besle et al., 2013; Ejaz et al., 2015; Gozzi et al., 2024; Kikkert et al., 2016; Kikkert et al., 2021; Kikkert et al., 2023; Kolasinski et al., 2016; Kuehn et al., 2018; Kuehn et al., 2014; Martuzzi et al., 2014; Odermatt et al., 2024; Puckett et al., 2017; Rabe et al., 2023; Sanchez-Panchuelo et al., 2012; Sanchez-Panchuelo et al., 2010; Sanders et al., 2023; Wesselink et al., 2019), several challenges arise when using it to assess somatosensory processing at the level of the brainstem (Beissner, 2015; Brooks et al., 2013; Sclocco et al., 2018). First, the brainstem is highly vulnerable to motion and physiological noise. Specifically, respiration and the pulsation of nearby arteries and cerebrospinal fluid (CSF) can introduce fMRI signal inhomogeneities, which severely impact signal quality (Brooks, Faull, Pattinson, & Jenkinson, 2013). Second, the somatosensory nuclei are small, measuring only a few millimetres in width and spanning just a few voxels in typical fMRI images. Moreover, the somatosensory nuclei, e.g. the cuneate and trigeminal, lay directly adjacent to each other, making it challenging to differentiate their activity given the spatial resolution limits of fMRI, which is typically 2-4mm (Glover, 2011; Menon & Goodyear, 2001). Finally, the brainstem serves as a primary hub for many diverse physiological processes, with nuclei responsible for autonomic control, pain modulation, and emotion processing closely surrounding those involved in sensorimotor signal transmission. This can increase uncertainty when attributing signals to individual nuclei (Harrison et al., 2023; Paxinos & Mai, 2004). Despite these limitations, recent advancements in fMRI acquisition and pre-processing techniques have significantly mitigated these challenges, enabling researchers to characterise brainstem activity in humans (Beissner et al., 2014; Harrison et al., 2023; Harvey et al., 2008; Matt et al., 2019; Reddy et al., 2024). However, very few fMRI investigations have explored human somatosensory brainstem processing.

In this study, we combined fMRI with a hand-and-face movement paradigm to map activation across the somatosensory nuclei of the brainstem, thalamus and S1. In comparison to sensory paradigms, motor tasks yield stronger and more robust univariate activity across the somatosensory stream (Charyasz et al., 2023; Komisaruk et al., 2002; Kündig et al., 2024; Sanders et al., 2023). We used both traditional univariate and state-of-the-art multivariate analysis approaches to assess the extent to which we can delineate hand and face activity across these somatosensory nuclei. If it is possible to distinguish the activity of adjacent nuclei with fMRI using univariate analysis, then we would expect a significant difference in hand- and face-task-specific activity across anatomical masks of the cuneate and spinal trigeminal nuclei. Importantly, multivariate classification approaches offer greater sensitivity for discriminating between subthreshold or overlapping representations (Ejaz et al., 2015; Haufe et al., 2014; Hebart & Baker, 2018; Muret et al., 2022; Wesselink et al., 2022). Therefore, we further anticipate that the multivariate classification analysis will provide improved differentiation of hand and face activity in the brainstem compared to the traditional univariate region-of-interest analysis. By integrating both univariate and multivariate analyses, we introduce a comprehensive methodological approach to explore signal processing in humans across the somatosensory stream.

## 2. Methods

### 2.1 Participants

We recruited 21 healthy adults between 18 and 75 years old. Given the novelty of this approach, there is no well-established method for calculating sample size. We conducted an a priori power analysis based on our previous study demonstrating significant S1 activity during hand movement (Kikkert et al., 2021), aiming for a power level of 80% for a two-tailed one-sample t-test (alpha level = 0.05), which resulted in a necessary sample size of 12 participants (calculated with G*Power; Faul et al., 2007) . However, group sizes larger than 20 are recommended for fMRI experiments and parametric statistical analysis (Pajula & Tohka, 2016), which informed our decision to recruit a larger sample size than required by the power analysis.

Exclusion criteria were as follows: no contraindications for MRI, absence of neurological disorders, no hand impairments, and the capacity to provide informed consent. Notably, one participant was rescanned due to abnormal fMRI noise components identified through independent component analysis (ICA). Another participant was excluded due to corruption of their fMRI data during data acquisition, resulting in a final sample of 20 participants (mean age ± s.e.m.= 50.8 ± 3.5 years; 2 females; 3 dominant left-handers). The study received ethical approval from the Ethics Committee of the Canton of Zürich (KEK-2018-00937), and written informed consent was obtained from all participants prior to inclusion. The study is registered on clinicaltrials.gov under NCT03772548.

### 2.2 Experimental design

We designed a blocked fMRI paradigm to investigate the activation of adjacent medullary nuclei involved in processing sensory input from the face and hands. The blocked design included three movement conditions: self-paced left-hand open-and-closing, right-hand open-and-closing, and face movement (i.e., continuous jaw clenching and self-paced lip pursing, as previously implemented by Beissner et al., 2011), as well as a rest condition. While tactile paradigms are an intuitive solution to localise sensory nuclei (Reddy et al., 2024; Komisaruk et al., 2002), motor tasks often evoke stronger and more robust activation across the somatosensory stream (Charyasz et al., 2023; Komisaruk et al., 2002; Sanders et al., 2023). This is particularly advantageous for imaging subcortical regions like the brainstem, which exhibit a lower signal-to-noise ratio (Brooks et al., 2013).

Instructions were presented as white text on a black screen (’left hand’, ’right hand’, ’face’, or ’rest’) and viewed by participants through a mirror mounted on the head coil. We further instructed participants to focus on a green fixation cross throughout the scan to minimise eye movements and avoid excessive activation of the abducens nucleus, located caudal to the principle sensory division of the trigeminal nuclei (Paxinos & Mai, 2004). Each movement block lasted 12 seconds, followed by a 6-second rest block, with each condition repeated eight times per run in a counterbalanced order (total run duration = 7min 12 sec). In total, we acquired four runs, each with a different block order, resulting in 32 trials per condition. Instructions were delivered using Psychtoolbox (Psychophysics Toolbox V.3), which was implemented in Matlab v9.5 (R2018b). Participant head motion was minimised by padded cushions and/or over-ear MRI-safe headphones.

### 2.3 MRI acquisition

We collected MRI data with a 3 Tesla Philips Ingenia system (Best, The Netherlands) and a 15-channel HeadSpine coil. We acquired a T1-weighted structural scan covering the brain and brainstem to facilitate the coregistration of functional images with anatomical regions of interest (sequence parameters: echo time (TE): 4.4ms; repetition time (TR): 9.3ms; flip angle: 8°; resolution: 0.75mm^3^). For task-fMRI, we implemented an echo-planar-imaging (EPI) sequence with partial brain coverage. We recorded blood-oxygenation-level dependent (BOLD) signals from 32 sagittal slices aligned parallel to the floor of the fourth ventricle, covering the brainstem, thalamus, and the primary somatosensory cortex (S1) (sequence parameters: single-shot gradient echo, TE: 26ms; TR: 2500ms; flip angle: 82°; spatial resolution: 1.45 x 1.8 x 1.45mm^3^; field of view (FOV): 210 x 187.6 x 70.2mm3; SENSE factor: 2.1; with 179 volumes per run). Additionally, we acquired a whole-brain, single-volume EPI image (with parameters identical to the task-fMRI protocol) to enhance co-registration between the fMRI data and anatomical scans. To facilitate post hoc physiological noise correction of the fMRI data, we recorded chest movement data using a respiratory bellow throughout the scans (sampling rate: 496Hz). A pulse oximeter was attached to the second digit of each participant’s foot to record oxygen saturation and cardiac pulse. All physiological measurement devices were connected to a data acquisition device (Invivo; Phillips Medical Systems, Orlando, Florida) coupled with a desktop computer running the corresponding recording software.

### 2.4 MRI analysis

Analysis of the MRI data was performed using tools from FSL v5.0.9 (http://fsl.fmrib.ox.ac.uk/fsl), Matlab v9.5 (R2018b), FreeSurfer v6.0 (https://surfer.nmr.mgh.harvard.edu/), SPM12 (http://www.fil.ion.ucl.ac.uk/spm/), the PhysIO toolbox (Kasper et al., 2017), the Decoding Toolbox (TDT) v3.999F (https://sites.google.com/site/tdtdecodingtoolbox/), and Connectome Workbench (https://www.humanconnectome.org/software/connectome-workbench).

#### 2.4.1 Preprocessing

As brainstem fMRI signals are highly susceptible to corruption by physiological and other noise sources, we applied an optimised brainstem fMRI preprocessing pipeline based on previous research (Beissner, 2015; Beissner et al., 2011; Faull et al., 2015; Matt et al., 2019; Meissner et al., 2024). This pipeline included brain extraction using FSL’s automated brain extraction tool (BET; Smith, 2002), motion correction using FMRIB’s Linear Image Registration Tool (MCFLIRT; Jenkinson et al., 2002), a 90s high-pass temporal filter implemented via FSL’s Expert Analysis Tool (FEAT; Woolrich et al., 2001), and spatial smoothing. We opted for a 2 mm full width at half maximum (FWHM) Gaussian kernel, as excessive smoothing of brainstem activity is not recommended due to a potential reduction in signal-to-noise ratio caused by averaging signal of interest with surrounding tissue. This kernel size is comparable to the diameter of the target nuclei and aligns with previous recommendations by Beissner et al. (2011).

Further denoising was completed using FSL’s Physiological Noise Modelling (PNM; Brooks et al., 2008). In this step, cardiac and respiratory phases were assigned slicewise to each volume of the EPI image time series. The physiological noise regression model comprised a total of 34 regressors: 4th-order harmonics for both the cardiac and respiratory cycles, 2nd- order harmonics for their interactions, and individual regressors for heart rate and respiration volume. We visually inspected the waveforms to ensure accurate identification of the peaks of the cardiac and respiratory cycles, making adjustments as needed. The resulting voxelwise confound regressors were then incorporated into the first-level general linear model (GLM).

For the multivariate analysis, physiological nuisance regressors were modelled as low-order Fourier series using the RETROICOR method (Glover et al., 2000), implemented in the PhysIO toolbox (Kasper et al., 2017) to allow compatibility with the first-level GLM in SPM12 (http://www.fil.ion.ucl.ac.uk/spm/). The regressors included 3rd-order cardiac harmonics, 4th- order respiratory harmonics, and their 1st-order interaction terms, along with individual regressors for respiratory volume per time (RVT) and heart rate variability, as recommended by Harvey et al. (2008).

Finally, independent component analysis (ICA) was conducted using the Multivariate Exploratory Linear Optimized Decomposition interface (MELODIC; Smith et al., 2004), which enables the decomposition of fMRI data into temporal and spatial components. The resulting components were classified as either “Noise” or “Signal” based on visual inspection of their spatial maps, time series, and power spectra information, following the guidelines published by Griffanti et al. (2017). Two independent researchers labelled the components to assess interrater reliability. Cohen’s kappa indicated strong agreement between raters, with an estimated score of 0.85 (95% CI: 0.82 to 0.88). The ICA-derived noise regressors were subsequently added to the GLM design matrix.

#### 2.4.2 Image Co-registration

Image co-registration was systematically completed in separate, visually inspected steps. First, we aligned each participant’s fMRI data to their whole-brain EPI image using FMRIB’s Linear Image Registration Tool (FLIRT), employing a mutual information cost function and 6 degrees of freedom (Jenkinson & Smith, 2001). Next, each participant’s EPI images were registered to the T1-weighted image, initially using 6 degrees of freedom and the mutual information cost function, and then optimised using boundary-based registration (BBR; Greve & Fischl, 2009). Subsequently, each T1-weighted structural image was co-registered to the MNI152 (1mm^3^) template using FMRIB’s nonlinear registration tool (FNIRT) with 12 degrees of freedom (Jenkinson, 2007). Following each step, the image co-registration was visually inspected using Freeview (FreeSurfer) to ensure precise alignment, particularly at the level of the medulla. In the instance of inaccurate registration between a participant’s fMRI data and T1-weighted image, co-registration of the fMRI data to the whole-brain EPI image was omitted, which was found to improve alignment at the level of the brainstem.

#### 2.4.3 Region of Interest Definition

To assess hand and face activity in our regions of interest (ROIs) along the somatosensory pathway, we defined masks of the brainstem, thalamus and postcentral gyrus in MNI 1mm^3^ space, as well as surface S1 hand and face area masks.

Whole masks of the right and left postcentral gyrus, bilateral thalamus and brainstem were derived from the Harvard-Oxford cortical and subcortical structural atlases (Desikan et al., 2006; Frazier et al., 2005). The superior limit of the postcentral gyrus mask was restricted to 1 cm above the anatomical hand knob in S1 (z = 63), and the inferior limit was set to the border of the secondary somatosensory cortex. The thalamus mask was thresholded at 10% to prevent overlap with the ventricles in the MNI 1mm^3^ brain template. Additionally, the brainstem mask was thresholded at 50%, and its caudal boundary was manually extended to include the full extent of the cuneate and spinal trigeminal nuclei (z = -71). To reduce the number of voxel-wise comparisons and increase the sensitivity of the univariate non-parametric group analysis, the brainstem mask was further restricted to include only the medulla by removing axial slices above the inferior pontine sulcus (z = -43). Notably, the whole brainstem mask was used for the multivariate classification analysis to avoid a strict a priori hypothesis regarding the location of decoding information and to exploit the covariation of data across the brainstem for noise suppression.

For the univariate ROI analysis, specific brainstem ROIs, including the spinal trigeminal (bilateral) and left and right cuneate nuclei, were manually defined on the MNI 1mm^3^ standard template brain. Notably, we selected the spinal nucleus of the trigeminal nerve as it runs immediately lateral to the cuneate nuclei, providing an ideal model for assessing our research question. We first determined the locations of the right cuneate and spinal trigeminal nuclei using anatomical coordinates extracted from a probabilistic stereotactic brainstem atlas published by Afshar (1978) and further detailed by Beissner et al. (2011). These masks were drawn relative to three anatomical planes: the fastigial floor line, the floor of the IVth ventricle, and the midsagittal plane. For each coordinate, the 90% confidence bounds were calculated and rounded to whole millimetres before drawing. To avoid overlapping ROIs, we excluded voxels that fell within the 90% confidence bounds of the spinal trigeminal and cuneate nuclei. The final volumes of the spinal trigeminal and the cuneate ROIs (unilateral) were 448 and 296mm^3^, respectively. Each ROI was then mirrored along the midsagittal plane to create the contralateral ROI. The cuneate masks were further restricted to below the level of the pyramidal decussation to limit potential signal overlap from both the left and right hands. Additionally, anatomic thalamic masks of the VPL and VPM nuclei were obtained from the Thalamus Morel Atlas (Krauth et al., 2010).

Finally, both S1 hand and face area ROIs were defined on the fs_LR template surface. The hand ROI was defined by selecting the nodes approximately ∼1 cm below and above the anatomical hand knob within S1, as defined by the Glasser et al. (2016) atlas. The face ROI was defined as starting 1cm lateral to the S1 hand area and included all S1 lateral to this boundary. A 1 cm gap was maintained between the S1 hand and face ROIs to exclude the portion of S1 where significant overlap between face and hand representations would be expected.

### 2.5 Univariate analysis

Time-series statistical analysis was performed per run using FEAT with FMRIB’s Improved Linear Model (FILM) with local autocorrelation correction (Woolrich et al., 2001). Activity estimates were derived from a general linear model (GLM), in which regressors were convolved with three basis functions generated from FMRIB’s Linear Optimal Basis Sets (FLOBS) toolkit (Mark W Woolrich et al., 2004). This approach was selected to accommodate the variability of BOLD responses in brainstem regions, which may not be adequately represented by the canonical HRF alone (Wall et al., 2009). While this method convolves the GLM regressors with both the canonical HRF and its temporal and dispersion derivatives, only the canonical HRF regression parameter estimates were used for subsequent higher-level analyses to prevent overestimating statistical effects (as implemented in Meissner et al. (2024) and Faull et al. (2015)). As detailed in Section 2.4.1, voxel-wise confound lists from PNM and ICA noise components were added to the GLM as nuisance regressors. Three main contrasts were defined for the analysis: each movement condition (left hand, right hand, face) versus rest. Additionally, the contrasts left-hand > all, right-hand > all, and face > all were defined for exploring whole-brain activity maps.

A second-level analysis was run for each participant to average across runs using FMRIB’s Local Analysis of Fixed Effects (Mark W. Woolrich et al., 2004). To minimise the risk of confounding results due to motion (e.g. noise-induced variability, partial volume effects, and motion-induced overlap of border voxels across ROIs), we excluded runs from the first-level analysis with an absolute mean displacement >1mm. In total, 4 out of the 80 acquired runs were excluded, with no more than one run being omitted per participant.

To visualise group-level activation, we conducted region-specific third-level statistical analyses utilising defined ROIs for the brainstem and thalamus as previously implemented in fMRI analyses of the subcortical regions (Brooks et al., 2017; Oliva et al., 2021; Oliva et al., 2022; Reddy et al., 2024). Notably, parametric, cluster-based correction methods, which are typically employed and suitable for widespread cortical activations, lack sensitivity to detect the smaller, more variable clusters in subcortical areas (Friston et al., 1994; Woo et al., 2014). Additionally, brainstem spatial smoothing constraints (as discussed in Section 2.4.1) are incompatible with parametric Gaussian field-based inferences, resulting in overly conservative significance estimates (Beissner et al., 2011; Worsley & Friston, 1995). Therefore, to increase sensitivity to activation in the brainstem and thalamus, we implemented FSL’s nonparametric permutation test RANDOMISE with threshold-free cluster enhancement (TFCE) and family-wise error (FWE) correction (Smith & Nichols, 2009; Winkler et al., 2014). Second-level COPE maps in 1mm^3^ MNI standard space were merged across participants for one-sample t-tests within anatomical masks of the medulla and thalamus (left and right combined) separately. The default parameters for variance smoothing (5mm; as in Reddy et al. (2024) and Kündig et al. (2024) and the number of permutations (5000) were utilised (Winkler et al., 2014). The resulting TFCE statistical p-maps were thresholded using a FWE correction threshold of p <0.05. However, as the complete activation patterns inform our multivariate approach, we utilise the ‘highlighting’ method proposed by Taylor et al. (2023) and additionally display subthreshold activity with a reduced opacity. For statistical inference of cortical activations, second-level statistical analyses were repeated in individual participant space. For optimal cortical alignment, the subsequent COPE maps were transformed to the cortical surface and concatenated for group-level analysis. The cortical surface projections were constructed from each participant’s native T1-weighted images. For each contrast of interest, we implemented FSL’s permutation analysis of linear models (PALM) with sign flipping and 5000 permutations of a two-sided one-sample t-test. The resulting group z-statistic images were thresholded at z > 3.1 and FWE corrected at p < 0.01.

### 2.6 Multivariate classification analysis

Multivariate classification is a machine learning technique that decodes different experimental conditions based on their neuronal activation patterns (Mur et al., 2009; Norman et al., 2006; Pereira et al., 2009). Unlike univariate ROI analyses, which provide insight into the strength of voxel-wise activations that are spatially averaged across a specified region, multivariate approaches evaluate simultaneous activity across multiple voxels. In the instance of overlapping activity, as anticipated in the brainstem, this approach may offer greater sensitivity for discriminating between conditions (Ejaz et al., 2015; Muret et al., 2022; Wesselink et al., 2022). Moreover, classification approaches incorporate subthreshold information to provide valuable insight into the underlying representational overlap (Haxby, 2012; Haxby et al., 2001; Haynes & Rees, 2006; Mur et al., 2009; Norman et al., 2006). While univariate approaches may demonstrate that two tasks activate the same voxel, they do not statistically evaluate this shared spatial representation, which is of key importance in assessing changes in somatotopic organisation. Importantly for our research question, multivariate analysis may provide a more comprehensive look at the extent to which we can separate between neural responses of adjacent brainstem nuclei. Therefore, we explored whether we could train a support vector machine (SVM) classifier to decode left-hand, right-hand and face activity within the brainstem, thalamus and S1.

For the classification analysis, we generated beta images for the three main contrasts (left-hand, right-hand and face versus rest) using a first-level GLM in SPM12. The settings were matched to those used in the univariate first-level GLM. We set the global masking threshold to zero to ensure that all voxels were retained for the classification analysis regardless of their intensity. As in the univariate analysis, we modelled both the temporal and dispersion derivatives and extracted the canonical HRF regression parameter estimates for classification. This resulted in a total of 240 beta images in 1mm^3^ MNI standard space, i.e. one beta per condition and per run for each participant. Transforming the first-level beta images to the standard template not only increases the number of samples to train the classifier but also allows us to classify across participants. Subsequently, we used the default model parameters of the decoding toolbox (TDT; Hebart et al., 2015) to implement a leave-one-participant-out cross-validation scheme (k = 20, L2-norm support vector machine with a linear kernel, C = 1). Notably, in contrast to traditional machine learning, the objective here is not to optimise the classifier’s performance but instead to demonstrate that the underlying brain signals contain condition-specific information relevant to classification. We calculated the decoding accuracy of each condition across all the voxels within masks of the brainstem, thalamus and S1 (see Section 2.4.3 for ROI details). We then extracted the average test classification accuracies and a confusion matrix of the predicted and true labels. Finally, we generated a null distribution from 1000 random permutations of the condition labels. We then calculated the empirical p-value of the average test classification accuracy based on the proportion of permutation accuracies equal to or greater than this score.

Importantly, the above classification analysis does not require a strict apriori hypothesis regarding the location of neural activity, unlike univariate ROI analyses, which are constrained by the need for precise ROI masks to capture activity accurately. Conversely, obtaining a significant classification accuracy across the brainstem does not provide insight into whether the classification analysis is sensitive to information encoded in the areas surrounding the target nuclei. Therefore, we also conducted a searchlight analysis to investigate which voxels within the brainstem were associated with classification accuracies above chance. We used a leave-one-participant-out cross-validation scheme (k = 20, L2-norm support vector machine with a linear kernel, C = 1). This analysis produces a searchlight map that assigns a multiclass classification accuracy to each voxel within the brainstem ROI. Notably, across-participant searchlight analyses assume that the spatial variation among participants is smaller than the searchlight radius and may otherwise be prone to spatial distortions or exaggeration (Etzel et al., 2013). For optimal sampling, Kriegeskorte et al. (2006) recommend a searchlight radius comparable to the size and compactness of the region. To balance these factors, we implemented a 3mm searchlight radius, which is comparable to the diameter of the targeted nuclei and may reduce the risk of spatial inaccuracies if the information is present in clusters within these nuclei (Afshar, 1978; Etzel et al., 2013; Knights et al., 2022).

To identify voxels with classification accuracies significantly above chance, we employed the statistical approach recommended by Stelzer et al. (2013). First, we ran 1000 permutations with random shuffling of the condition labels to create a null distribution of classification accuracies across all voxels in the brainstem. We then determined the classification accuracy corresponding to a critical cut-off of 0.05, thresholding both the initial searchlight map and the 1000 permutation maps by this value. Next, we conducted cluster correction by extracting the maximum cluster size within the thresholded permutation maps to create a null distribution of maximum cluster sizes. Finally, we calculated the cluster size corresponding to a p-value of 0.05 and masked the thresholded searchlight map to only include clusters larger than this value.

### 2.7 Statistical analysis

Statistical analysis for the univariate ROI analysis was conducted using RStudio (v4.0.3) and GraphPad Prism (v10.3.0). Outlier values ± 2.5 standard deviations from the mean were replaced with the mean. All statistical testing was conducted with and without outliers to confirm that this adjustment did not change the results. We used standard approaches for statistical analysis, as detailed in the Results section. Parametric or non-parametric testing was used depending on normality, which was confirmed with the Shapiro-Wilk test. In addition, Mauchly’s test was implemented to assess sphericity across each ROI, and Bonferroni or Dunn’s (Friedmans ANOVA) corrections were applied for post-hoc comparisons. Planned contrasts were implemented to compare the mean of the target condition with each non-target condition.

## 3. Results

We investigated whether 3T fMRI, with an optimised acquisition and preprocessing pipeline, could effectively distinguish between neural responses of neighbouring somatosensory nuclei in the brainstem, thalamus and S1. We delineated left-hand, right-hand, and face movement-related activity utilising two approaches. First, using traditional univariate voxel-wise and ROI analyses, we assessed the extent to which we could distinguish between conditions based on their overall activation strength. Second, using multivariate classification analysis, we explored the degree to which somatosensory nuclei can be distinguished based on their underlying activation patterns for the different conditions, thereby providing greater insight into the representational overlap of these signals. In addition, we conducted a multiclass searchlight analysis to determine the specific regions of the brainstem encoding relevant information for the classification of hand and face movement.

### 3.1 Univariate analysis reveals a partial differentiation of hand and face activation within somatosensory relay nuclei

We first explored body-part-specific activity across the somatosensory processing stream. To do so, we created group activity maps for each condition contrasted with all other conditions. We found significant group-level activation in the anticipated nuclei and areas of the brainstem, thalamus, and S1 (Figure 2a).

**Figure 2.**
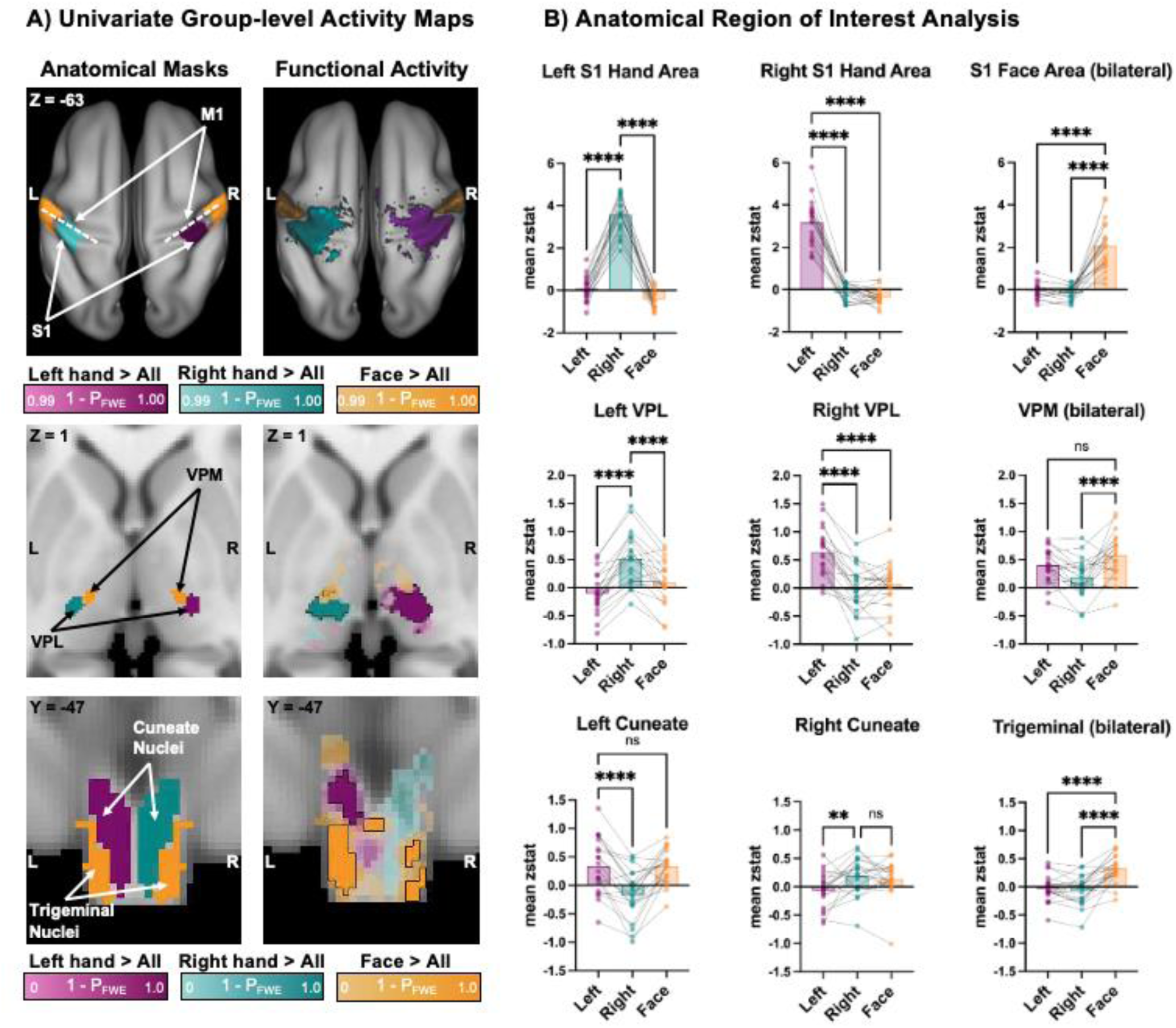
Univariate analyses can partially differentiate hand and face activity within neighbouring somatosensory nuclei. (A) Group-level activity maps (n = 20) during left-hand, right-hand and face movements. To obtain body-part-specific activity, we contrasted the conditions as follows: left-hand > right-hand and face (purple), right-hand > left-hand and face (turquoise), and face > left and right hands (yellow). Anatomical regions of interest for the somatosensory processing stream are displayed in the left column, demonstrating anticipated areas of activation (left cuneate nucleus = purple; right cuneate nucleus = turquoise; spinal trigeminal nucleus = yellow). The corresponding family-wise error-corrected (P_FWE_) activity maps are shown in the right column. Cortical activations are projected onto a cortical surface, thresholded at Z > 3.1 and P_FWE_ < 0.01; subthreshold cortical activity is omitted for clarity. For the brainstem and thalamus, suprathreshold activity clusters are outlined in black (threshold-free cluster enhancement, P_FWE_ < 0.05). Subthreshold activity is shown with reduced opacity. P_FWE_ values are indicated alongside their respective colour bars. (B) Region of interest (ROI) analysis shows the mean z-statistic for each task (contrasted with rest) within anatomical masks of the left cuneate, right cuneate and spinal trigeminal nuclei (visualised in A, left panel). Individual participants are plotted as dots. Statistical significance is denoted by: **** = p < .0001, *** = p < .001, ** = p < .01. P-values are corrected for multiple comparisons within each ROI.

At the cortical level, we found significant task-selective brain activity during left- and right-hand movement in the contralateral S1 and primary motor (M1) hand cortices (Figure 2a; top panel). We further detected significant bilateral activity elicited by face movement laterally to the hand representations, consistent with the established somatotopic organisation of these body parts in the cortex (Penfield & Boldrey, 1937; Penfield & Rasmussen, 1950; Roux et al., 2018). To enhance sensitivity for detecting subcortical activation, we conducted region-based group-level analyses within anatomically defined masks of the medulla and thalamus. As expected, left- and right-hand movements significantly activated the contralateral VPL nucleus of the thalamus (Figure 2a; middle panel). In contrast, face movement did not yield significant activation in the right thalamus. However, uncorrected TFCE p-maps revealed subthreshold face-task-selective activation medial to the hand clusters, overlapping with the VPM nuclei. In the medulla, left-hand movement elicited significant activation clusters overlapping with the left cuneate nucleus. While the right-hand movement did not produce significant clusters, subthreshold activation was localised within the right cuneate nucleus (Figure 2a; bottom panel). Face movements significantly activated the spinal trigeminal nuclei bilaterally, as expected. Notably, this activation was widespread, with one significant cluster located between the cuneate nuclei.

Next, we aimed to explore the extent to which univariate activity can be distinguished within precise ROIs of the somatosensory relay nuclei. To do so, we extracted the task-related activity, quantified as the mean z-statistic, for each movement condition compared to rest within anatomical masks of the cuneate and spinal trigeminal nuclei (brainstem), the VPL and VPM (thalamus), hand and face cortical areas (S1) (Figure 2b). During movement of the hands, we anticipated higher activation in the ipsilateral cuneate, contralateral VPL and contralateral S1 hand area, as compared to the opposite hand and face movement. Conversely, during movement of the face, we anticipated higher bilateral activation within the spinal trigeminal, VPM and S1 face areas, as compared to left or right-hand movement.

The statistical results of the ROI-specific comparisons are summarised in Table 1. In S1 (Figure 2b; top panel), we reliably distinguished between the hand and face movement conditions across all ROIs. Similarly, in the thalamus (Figure 2b; middle panel), we were able to distinguish between hand and face movement within the left and right VPL ROIs. However, in the VPM ROI, we only found a trend towards higher face than left-hand activity. Finally, in the brainstem (Figure 2b; bottom panel), while we could distinguish between ipsilateral and contralateral hand activity in the cuneate ROIs, we could not distinguish between the ipsilateral hand and adjacent face activity. These findings suggest a limited ability to differentiate between hand and facial activity within the anatomical confines of our cuneate ROIs using univariate analysis. However, in support of our hypothesis, we found that face activity was significantly higher than both left- and right-hand activity within the spinal trigeminal ROI.

**Table 1.** Summary of statistical effects for the univariate ROI analyses.

| ROI | comparison | Test Statistic | p-value | Effect size |
| --- | --- | --- | --- | --- |
| *S1* |  |  |  |  |
| **Left S1 Hand Area** | Main effect | $F(2, 38) = 303.4$ | $< .0001$ | $\eta_p^2 = 0.941$ |
| | Right vs. Left hand | $t(38) = 19.74$ | $< .0001$ | |
| | Right hand vs. Face | $t(38) = 22.63$ | $< .0001$ | |
| **Right S1 Hand Area** | Main effect | $\chi^2(2) = 30.40$ | $< .0001$ | $W = 0.760$ |
| | Left vs. Right hand | $Z = 4.427$ | $< .0001$ | |
| | Left hand vs. Face | $Z = 5.060$ | $< .0001$ | |
| **Bilateral S1 Face Area** | Main effect | $F(2, 38) = 54.64$ | $< .0001$ | $\eta_p^2 = 0.742$ |
| | Face vs. Left hand | $t(38) = 7.459$ | $< .0001$ | |
| | Face vs. Right hand | $t(38) = 7.446$ | $< .0001$ | |
| *Thalamus* |  |  |  |  |
| **Left VPL** | Main effect | $F(2, 38) = 52.63$ | $< .0001$ | $\eta_p^2 = 0.735$ |
| | Right vs. Left hand | $t(38) = 10.08$ | $< .0001$ | |
| | Right hand vs. Face | $t(38) = 6.687$ | $< .0001$ | |
| **Right VPL** | Main effect | $F(2, 38) = 36.13$ | $< .0001$ | $\eta_p^2 = 0.655$ |
| | Left vs. Right hand | $t(38) = 7.841$ | $< .0001$ | |
| | Left hand vs. Face | $t(38) = 6.763$ | $< .0001$ | |
| **Bilateral VPM** | Main effect | $F(2, 38) = 10.54$ | $< .001$ | $\eta_p^2 = 0.357$ |
| | Face vs. Right hand | $t(38) = 4.582$ | $< .0001$ | |
| | Face vs. Left hand | $t(38) = 2.052$ | $0.094$ | |
| *Brainstem* |  |  |  |  |
| **Left Cuneate** | Main effect | $F(2, 38) = 24.40$ | $< .0001$ | $\eta_p^2 = 0.562$ |
| | Left vs. Right hand | $t(38) = 6.079$ | $< .0001$ | |
| | Left hand vs. Face | $t(38) = 0.061$ | $> .9999$ | |
| **Right Cuneate** | Main effect | $\chi^2(2) = 9.70$ | $0.008$ | $W = 0.243$ |
| | Right vs. Left hand | $Z = 3.004$ | $0.0053$ | |
| | Right hand vs. Face | $Z = 0.791$ | $0.858$ | |
| **Spinal Trigeminal** | Main effect | $F(2, 38) = 32.25$ | $< .0001$ | $\eta_p^2 = 0.629$ |
| | Face vs. Left hand | $t(38) = 6.708$ | $< .0001$ | |
| | Face vs. Right hand | $t(38) = 7.179$ | $< .0001$ | |

Overall, these results indicate that activation among somatosensory nuclei can be distinguished to some extent using univariate analysis of 3T fMRI data. However, this distinction varies across conditions and locations. Specifically, our brainstem ROI analysis of the cuneate nuclei did not clearly differentiate between hand and facial activation. The anatomical definition of our ROIs was designed so that the cuneate and spinal trigeminal nuclei were adjacent, with no intervening gap. This approach is constrained by the need for precise ROI masks to accurately capture activity. Therefore, to examine the extent to which we can distinguish between conditions without being limited by this potential confound, we next conducted a multivariate classification analysis.

### 3.2 Multivariate classification can differentiate between the activation of neighbouring brainstem nuclei involved in somatosensory processing

To investigate the extent to which left-hand, right-hand and face movement can be distinguished through their neuronal activity patterns, we conducted a classification analysis. Specifically, we tested whether we could train a support vector machine (SVM) to decode left-hand > rest, right-hand > rest and face > rest activation across participants within anatomical masks of the brainstem, thalamus and S1. As anticipated, we found high classification accuracy in S1, where fMRI signals are more robust and less prone to noise corruption, with an average classification accuracy across participants of 98.3% (p = <.001; empirical chance level: 33.3%). Moreover, a confusion matrix comparing the frequency of predicted to true labels did not display a bias towards an individual movement condition (Fig. 3). Importantly, we found that condition-specific activity could also be distinguished significantly above chance level in both the thalamus (accuracy = 60.4%, p = <.001) and the brainstem (accuracy = 50.8%, p = <.001) (Fig. 3). The confusion matrices revealed that we could distinguish between conditions in both these subcortical areas. These results indicate that there is sufficient information present in 3T fMRI data subcortically to decode between somatosensory representations of the face and hands.

**Figure 3.**
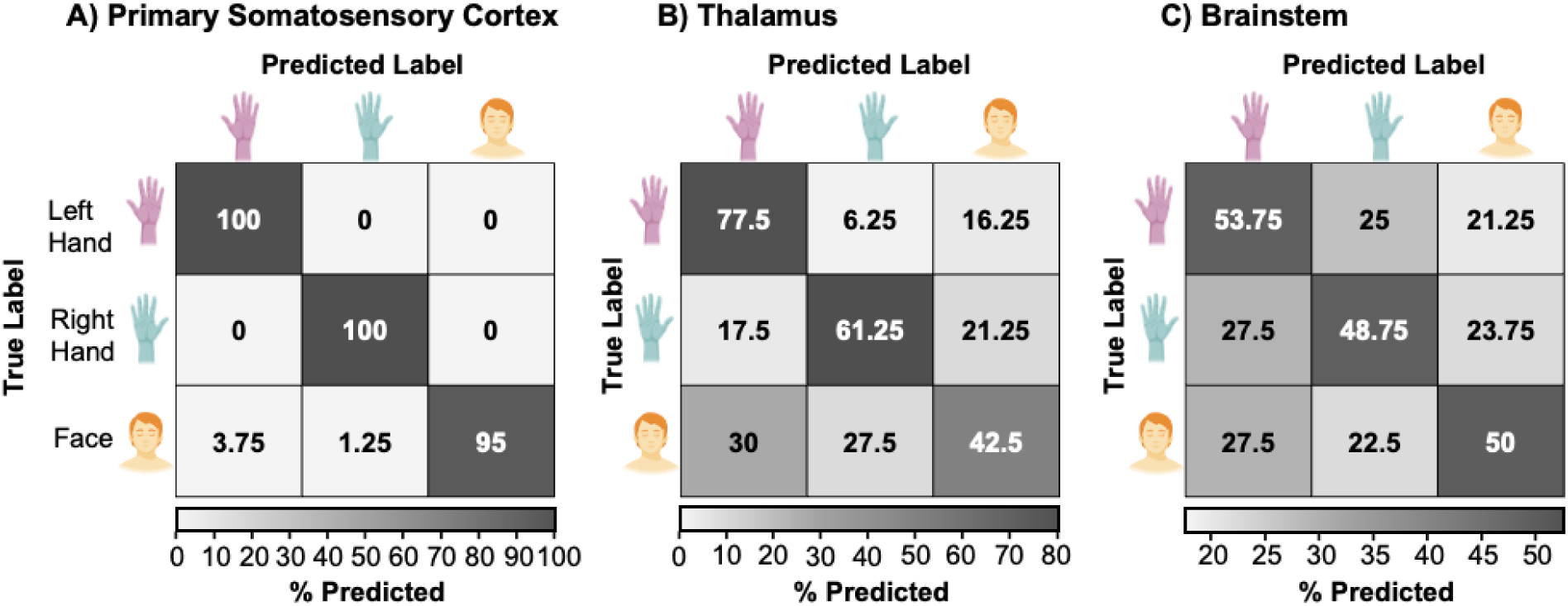
Hand and face activation patterns can be distinguished across the somatosensory processing stream using multivariate classification analysis. We trained linear support vector machines to classify voxel-wise activity patterns across participants in three anatomical regions of interest (ROIs): (A) brainstem, (B) thalamus, and (C) primary somatosensory cortex (S1). Each confusion matrix depicts the percentage of correct and incorrect classifications for left hand > rest, right hand > rest, and face > rest contrasts (rows = true labels, columns = predicted labels).

While the classification analysis described above shows that we can reliably decode between conditions based on brainstem activity patterns, it does not specify which regions of the brainstem enable our classifier to differentiate between conditions. We, therefore, conducted a searchlight analysis to investigate which voxels within the brainstem were associated with classification accuracies above chance level. The resulting accuracy map (Fig. 4) was thresholded by a critical cut-off of 39.7% accuracy (p < .05) and cluster-corrected (k = 639 voxels; p < .05). As anticipated, the voxels with significant classification accuracies were localised in the dorsal medulla, overlapping with the anatomical masks of our target nuclei: the cuneate and spinal trigeminal nuclei. More extensive significant clusters were observed in the left dorsal medulla. Significant voxels were also located in the dorsal pons surrounding the trigeminal motor and principal sensory nuclei. This result demonstrates that using multivariate classification analysis, 3T fMRI, and a body movement task, it is possible to distinguish between the neuronal activity patterns of adjacent brainstem nuclei involved in somatosensory processing.

**Figure 4.**
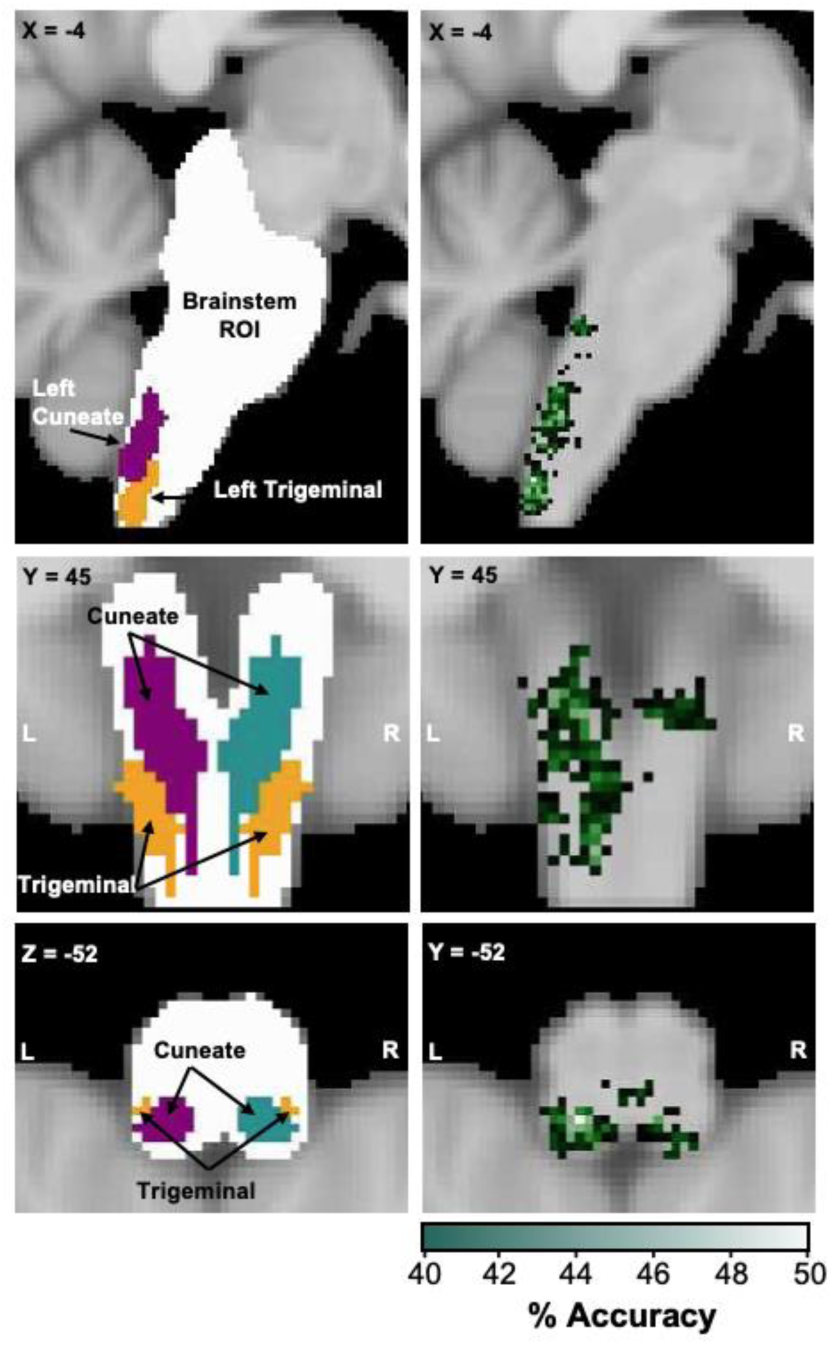
Searchlight analysis across the brainstem demonstrates that voxels differentiating between face and hand movement are localised in the dorsal medulla. We conducted a searchlight analysis to identify the contribution of individual brainstem voxels to the classification of hand and face movements. The left panel demonstrates the brainstem region of interest (white) implemented in the analysis. The expected areas of neural activation for somatosensory input from the face (spinal trigeminal nucleus; yellow) and hands (cuneate nucleus; purple) are overlaid. The right panel displays the searchlight accuracy map illustrating voxels with classification accuracies significantly above chance (thresholded and cluster-corrected at a critical cut-off of 39.7%, p = < .05; k = 639 voxels). The results indicate that the activation patterns for hand and face movement are predominately located in the dorsal medulla, overlapping with the anticipated nuclei involved in the somatosensory processing of face and hand input.

## 4. Discussion

Functional mapping of the human brainstem presents many challenges. These methodological constraints have deterred essential investigations into human sensory brainstem processing. Here, we delineate the functional activity of two adjacent somatosensory nuclei, namely the spinal trigeminal and cuneate nuclei. Our results demonstrate that functional activity can be distinguished in humans across the somatosensory stream through fMRI and optimised data acquisition and analysis strategies. This is a fundamental step towards broadening our understanding of somatosensory processing within humans and determining what changes in organisation may occur following sensory deprivation.

To date, human imaging studies involving the cuneate and trigeminal nuclei have primarily demonstrated activation in these regions as secondary outcomes whilst investigating alternate physiological processes (Faull et al., 2015; Pattinson et al., 2009) or while optimising brainstem pre-processing methods (Beissner et al., 2011; Harvey et al., 2008). Crucially, researchers have yet to establish the extent to which fMRI can be used to delineate the functional activity of these neighbouring nuclei. Notably, previous studies have demonstrated the capability of fMRI to localise univariate activity in adjacent sensory and motor brainstem nuclei (Komisaruk et al., 2002; Reddy et al., 2024). However, these studies did not directly delineate the extent to which these overlapping activations could be differentiated. This distinction between localisation and discrimination is crucial for determining the somatotopic organisation of somatosensory representations and assessing if activity is reorganised in response to environmental stimuli or injury.

We utilised both univariate and multivariate analyses to delineate the activity of neighbouring somatosensory nuclei. Notably, in the univariate group analysis, neither right-hand activity clusters in the brainstem nor face activity clusters in the right thalamus reached the significance threshold. A previous study by Reddy et al. (2024) demonstrated significant activation in the brainstem during both hand and foot stimulation using fMRI. However, while Reddy et al. contrasted each body part with rest, we compared each body part to all other conditions. Therefore, instead of simply identifying clusters associated with each body part, we sought body-part selective clusters to assess how effectively we could differentiate between tasks. Importantly, the uncorrected group activity maps revealed that, although weak, face and right-hand selective clusters spatially overlapped with the anticipated nuclei involved in somatosensory processing. This finding highlights a limitation of univariate analyses, which are constrained by the chosen significance threshold, whereas multivariate analyses can combine subthreshold information across voxels to enhance signals that may otherwise go undetected (Hebart & Baker, 2018).

Similarly, in the univariate ROI analysis, while we were able to distinguish between all conditions in S1, we were only able to partially differentiate between conditions in the brainstem and thalamus. Given that atlas-based masks do not draw a distinct border between the hand and face nuclei in these regions, even slight anatomical differences between individuals could cause inaccuracies in estimating these borders, leaving no margin for error. Moreover, the normalisation process during coregistration may have caused a slight misalignment of the anatomical borders in individual participants, causing an artificial overlap in activity. As with the S1 analysis, it would be beneficial to leave a border of empty voxels between ROIs to minimise this potential overlap. However, the small size of the brainstem nuclei and the resolution limits of 3T imaging make such an approach unfeasible. Higher-field imaging, such as 7T fMRI, may provide the spatial resolution needed to define a precise boundary between nuclei. However, it may also increase susceptibility to artefacts and signal inhomogeneities (Smith et al., 2014). Alternatively, the inability to separate between hand and face activity may reflect a (partially) shared somatotopic overlap between these representations within the cuneate nucleus, as previously observed in the human primary sensorimotor cortex (Gordon et al., 2023; Muret & Makin, 2021; Muret et al., 2022; Schellekens et al., 2018). Indeed, separation between the two nuclei has proved unsuccessful in animal research, where direct injection of tracers has led to labelling of the neighbouring nucleus (Jain et al., 2000; Liao et al., 2021; Marfurt & Rajchert, 1991). Therefore, analysis techniques that do not rely on such a strict a priori region specification may be more appropriate for differentiating between these potentially overlapping somatosensory representations.

Multivariate classification analyses provide a solution to evaluating neuronal representations across conditions within the entire somatosensory system. Notably, this technique offers greater sensitivity and specificity than univariate analysis by targeting distributed patterns of activity and exploiting the covariance of data across voxels (Haynes & Rees, 2005; Hebart & Baker, 2018; Jimura & Poldrack, 2012; Yukiyasu Kamitani & Frank Tong, 2005). More importantly, this method may provide a more comprehensive understanding of how well we can separate neural responses of adjacent brainstem nuclei, even when the signal-to-noise ratio is low (Haufe et al., 2014; Hebart & Baker, 2018). Indeed, through this approach, we found that we could differentiate between the neural representations of left-hand, right-hand and face movement across the brainstem, thalamus and S1. To infer which areas of the brainstem encoded informational content relating to these tasks, we also conducted a searchlight analysis. We found that the hand and face movement conditions were specifically encoded in the dorsal medulla, overlapping with the anticipated nuclei involved in somatosensory processing. To our knowledge, multivariate classification has not yet been applied to distinguish body-part selective activity within the brainstem. Our findings demonstrate a novel solution for exploring signal processing within the brainstem and across the entire somatosensory system. Moreover, given the brainstem’s diverse roles in pain modulation (Youssef et al., 2016), auditory processing (De Martino et al., 2013), visual perception (Limbrick-Oldfield et al., 2012), respiration (Faull et al., 2015), emotion processing (Satpute et al., 2013), behavioural arousal (Aston-Jones et al., 1999), and autonomic control (Samuels & Szabadi, 2008), we suggest that multivariate approaches may allow crucial insights into a broad range of physiological processes.

Importantly, in our classification analysis, we utilised an across-participant decoding design as opposed to the approach that is commonly used for cortical classification analysis, where inferences are made by combining within-participant accuracy scores (Gallivan et al., 2011; Haxby et al., 2001; Y. Kamitani & F. Tong, 2005; Nambu et al., 2015; Odermatt et al., 2024). In regions with a low SNR, like the brainstem, aggregating data across participants may provide more robust results and has been demonstrated to detect smaller effects if the inter-individual variability remains moderate (Wang et al., 2020). Critically, this method enables us to make inferences about somatosensory processing differences between patient versus control populations. Therefore, future studies may extend this analysis to investigate changes in somatotopic organisation between healthy and patient cohorts that experience sensory deprivation, e.g., spinal cord injury, limb loss, or peripheral neuropathy. Understanding the extent to which neural representations of sensory stimuli are preserved or altered following major sensory deprivation may assist in the development of therapeutic interventions that aim to promote adaptive plasticity and prevent the development of maladaptive circuitry.

Together, our results demonstrate that sensory activation in adjacent brainstem nuclei can be distinguished in healthy individuals using a movement paradigm and 3T fMRI with optimised data acquisition and analysis strategies. This provides the methodological foundation for future investigations into a wide range of brainstem-mediated functions, including the plasticity or stability of somatosensory processing in clinical populations.

## Acknowledgements

We thank the participants for their participation in the study. We thank Sijamini Baskaralingam and Charlotte Meninghin for their help with data collection. We thank Roger Luchinger and the Institute for Biomedical Imaging for scanning support. This project, P. Howell and S. Kikkert are funded by the Swiss National Science Foundation Ambizione Grant PZ00P3_208996 and Grant 32003B_207719. N. Wenderoth is additionally supported by the National Research Foundation, Prime Minister’s Office, Singapore under its Campus for Research Excellence and Technological Enterprise (CREATE) program (FHT). O. Harrison is supported by the Royal Society of New Zealand and New Zealand Lottery Health Research. S.N. Meissner is supported by the ETH Career Seed Award.

